# PanCNV-Explorer: Deciphering copy number alterations across human cancers

**DOI:** 10.64898/2026.01.26.701884

**Authors:** Nadine S. Kurz, Kevin Kornrumpf, Anna-Rosa Krüger, Jürgen Dönitz

## Abstract

Copy number variants (CNVs) are major drivers of cancer progression and genetic disorders, yet their interpretation, spanning biological mechanisms, clinical relevance, and therapeutic implications, remains fragmented across disparate resources. To bridge this gap, we present PanCNV-Explorer, a unified database integrating harmonized copy number variation data across 33 cancer types, cancer cell lines, and healthy cohorts. PanCNV-Explorer provides a genome-wide atlas of CNV frequency and functional impact, quantifying tissue-specific amplifications and deletions in both cancer and non-cancer contexts through rigorous cross-dataset normalization. The interactive web interface enables researchers to dynamically query CNVs by genomic coordinates or gene symbol, visualize cancer-type-specific frequencies with real-time comparative analysis, and explore integrated genomic features including transcripts, regulatory elements, and gene expression through a zoomable genome browser. Beyond exploration, the platform offers programmatic APIs for pan-cancer CNV analysis and visualization. A public web instance of PanCNV-Explorer is available at https://mtb.bioinf.med.uni-goettingen.de/pancnv-explorer/.

## 1 Introduction

Copy number variations (CNVs) are structural genetic alterations involving gains or losses of DNA segments that have profound implications for human health and disease [1]. Detected in over 40% of tumors [2, 3], CNVs directly dysregulate oncogenes and tumor suppressors through dosage effects [4] and are established as major drivers of cancer [5, 6, 7, 8]. Pan-cancer analyses reveal striking cancer-type specificity: Recurrent alterations occur in distinct genomic regions across cancer types [9], and mutational signatures exhibit significant inter-tumor heterogeneity [7]. This heterogeneity underscores the need for cancer-type-specific interpretation of somatic CNVs. Clinically, somatic CNVs guide targeted therapy, such as *ERBB2* amplifications predicting trastuzumab response in breast cancer. They also define molecular subtypes; for example, *CDKN2A/B* deletions characterize the activated B-cell-like (ABC) subtype of diffuse large B-cell lymphoma (DLBCL) [10]. Critically, distinguishing pathogenic somatic CNVs from benign germline variants or tissue-specific polymorphisms is essential for accurate clinical interpretation.

GISTIC2.0 [11] remains the community standard for identifying recurrent somatic CNVs within single cohorts, but lacks robust statistical methods for cross-cancer comparison of CNV frequencies due to cohort size disparities and batch effects. Similarly, tools like GATK-gCNV [12] and CNVkit [13] can distinguish somatic from germline CNVs using matched normal samples, yet lack cancer-type-specific baselines to differentiate somatic drivers from benign tissuespecific germline variants. Furthermore, annotation pipelines (e.g., AnnotSV [14], StrVCTVRE [15]) and pathogenicity predictors [16, 17, 18, 19, 20] primarily assess individual variants in isolation rather than integrating cancer-specific patterns or functional impacts. Existing databases (e.g., 1000Genomes, gnomAD, dbVar, DGV) catalog germline copy number variation, but do not provide cancer-specific somatic CNV data. Although resources like CNVIntegrate [21] and CNAapp [22] offer partial solutions, they lack multi-ethnic baselines, region-specific significance testing, or statistical frameworks for cross-cancer frequency comparisons. Critically, while integrative pan-cancer analysis of CNV-functional consequence correlations (e.g., gene expression dysregulation) holds significant promise [23], no publicly available tool currently enables this approach.

To address these gaps, we introduce PanCNV-Explorer, a harmonized database integrating somatic CNVs from 33 cancer types with germline CNVs from multi-ethnic healthy populations, dynamic cross-cancer comparison frameworks, or region-specific significance testing. Our platform provides an interactive web interface for real-time exploration of CNV landscapes with cross-cancer statistical validation using cohort-normalized metrics. Unlike recurrence-based approaches, PanCNV-Explorer distinguishes CNV drivers from passengers by identifying significant deviations from population-specific healthy baselines, thereby enhancing biomarker discovery and reducing false positives from benign variants.

## 2 Methods

### 2.1 Datasets

To retrieve cancer-specific CNVs, we downloaded copy number segmentation files from The Cancer Genome Atlas (TCGA) [24] from 33 cancer types using the TCGAbiolinks package [25]. To ensure sample uniqueness and consistency with contemporary best practices, we restricted analysis to samples processed with GATK4. Both TCGA samples with a segment mean <-0.2 were classified as deletions, and those with >0.2 as amplifications, aligning with thresholds that have been employed in prior works [26].

For cancer cell line data, we downloaded segment-level copy number variation data from DepMap (release 25Q3). The DepMap dataset was log_2_-transformed, and identical thresholds as those used for the TCGA data were applied. To characterize CNVs for healthy and normal tissue, we downloaded the structural variants calls of dbVAR from https://ftp.ncbi.nlm.nih.gov/pub/dbVar/data/Homo_sapiens/by_assembly/GRCh38/vcf/GRCh38.variant_call.all.vcf.gz. Non-human entries and aggregated population-level calls were excluded during preprocessing. Pan-tissue CNV data from DGV was downloaded in GRCh38 (2020-02-25 release) [29]. To derive copy number variants and structural variants from the gnomAD database (v4) [30], we downloaded non-neuro controls structural variants from https://storage.googleapis.com/gcp-public-data--gnomad/release/4.1/genome_sv/gnomad.v4.1.sv.non_neuro_controls.sites.vcf.gz and exome CNV non neuro controls from https://storage.googleapis.com/gcp-public-data--gnomad/release/4.1/exome_cnv/gnomad.v4.1.cnv.non_neuro_controls.vcf.gz. To standardize annotation and analysis, all CNV files were converted to BED format.

### 2.2 Data harmonization

To harmonize the cancer types between the TCGA and Depmap datasets, we created a manual mapping of the DepMap entities (*DepMapModelType*) to the TCGA tumor types. DepMap cancer types that could not be mapped to the TCGA classification were not included in the dataset. For dbVar, DGV, and gnomAD, we retained the original variant classifications (deletion/duplication) provided in the source files. Duplications were merged based on identical genomic coordinates (`chrom, start, end`) and sample identifiers.

To harmonize CNV breakpoints across datasets, we computed consensus segments using a three-step approach: For each chromosome *c*, we collected all unique start (*s*_*i*_) and end (*e*_*i*_) posi-tions from CNV calls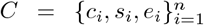,forming the breakpoint set *B* Sorting *B*_*c*_ yields *B*_*c*_ = *b*_1_ < *b*_2_… < *b*_*m*_, and the consensus segments are: *S*_*c*_ = (*b*_*j*_, *b*_*j*+1_ *−*1)| *j* = 1, …, *m −*1 We then applied breakpoint clustering with parameter *δ*, where break-points were grouped into clusters where consecutive points differ by ≤ *δ*, then replaced each cluster with its median. The resulting segment set spanned 200.000 segments throughout the whole genome. Fixed-sized bins divided the whole genome into segments of 10.000 base pairs per segment. Gene start and end positions were retrieved by extracting protein-coding genes from GENCODE v49 [31].

The frequency of a CNV in a specific tissue type was determined by calculating the overlaps between CNV calls and our consensus segments across all cancer types and healthy tissues, separately for amplifications and deletions. For individual-sample datasets (TCGA, DepMap), we counted samples with CNVs overlapping each segment. For aggregated population databases (gnomAD, DGV, dbVar), we extracted precomputed allele frequencies and remapped them to our unified segments using a binary overlap approach that preserves the original sample resolution.

Let *s*∈ *S* represent a unified genomic segment with coordinates (*c*_*s*_, *b*_*s*_, *e*_*s*_), where *c*_*s*_ is the chromosome identifier, and [*b*_*s*_, *e*_*s*_) denotes the half-open genomic interval. For each tissue type *t* ∈𝒯 and mutation type ℳ ∈ ℳ (where ℳ = {amplification, deletion}), we calculated the set of CNVs overlapping segment *s* as:

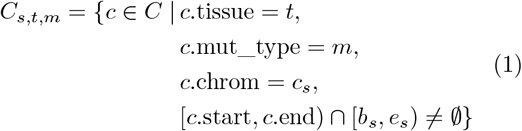

The total number of samples with CNVs overlapping segment *s* in tissue *t* for mutation type *m* is:

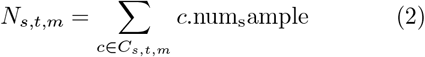

where c.num_sample represents either the count of samples carrying CNV *c* (for aggregated databases), or 1 for each individual sample (for non-aggregated datasets like TCGA). The CNV frequency for segment *s* in tissue *t* is then calculated as:

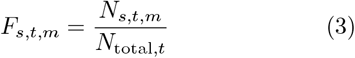

where *N*_*total,t*_ is the total number of samples in tissue *t*. This methodology preserves the original resolution of both individual-sample and aggregated datasets while enabling direct comparison across diverse resources. For gene-level analysis, CNV frequencies required >50% reciprocal overlap between the CNV and the gene body.

### 2.3 Database construction

The CNV database was created using PostgreSQL by generating the database from the original table-formatted files with the SQLAlchemy package [32].

### 2.4 The web server and front end

The database was built using PostgreSQL v18.1. The PanCNV-Explorer web server was built using Python Flask. The web front end for PanCNV-Explorer was implemented using AngularJS [33]. The genome browser tracks were implemented using JBrowse2 [34]. Interactive visualizations were generated using Plotly [35]. API documentation was realized using Swagger UI. All modules, including the database, server and front end, were containerized using Docker [36].

### 2.5 CNV annotation modules

To annotate copy number alterations, we created multiple web services providing CNV annotation for a variety of features. Affected genes of CNV regions were retrieved using the SeqCAT [37]. Data on gene dosage sensitivity, including haploinsufficiency (HI) and triplosensitivity (TS) scores, were retrieved from the ClinGen database [38]. Oncogenes and tumor suppressor genes classifications were downloaded from the Network of Cancer Genes database (NCG 7.0) [39]. To identify CNVs in API requests, we used the annotation <chr>:<start>-<end><change>, e.g. chr1:258946-714338_DEL. To provide annotations of the pathogenicity of CNVs, we developed annotation modules based on the methods CNVoyant [16], X-CNV [17], dbCNV [18], ISV[40], TADA [19], ClassifyCNV [20].

## 3 Results

### 3.1 A harmonized resource for somatic and germline CNV analysis

PanCNV-Explorer provides a comprehensive catalog of somatic copy number variations in cancer genomes and germline CNVs in normal human populations across diverse tissues. We integrated segment-level CNV data from tumor samples across 33 cancer types in TCGA and DepMap, alongside germline CNV data from 1000 Genomes, gnomAD, DGV, and dbVar (encompassing >100,000 individuals). Following classification of TCGA/DepMap CNVs as somatic events (Figure 1), we harmonized all data to a common BED format. To enable scalable analysis, we generated three complementary database versions:

**Figure 1:**
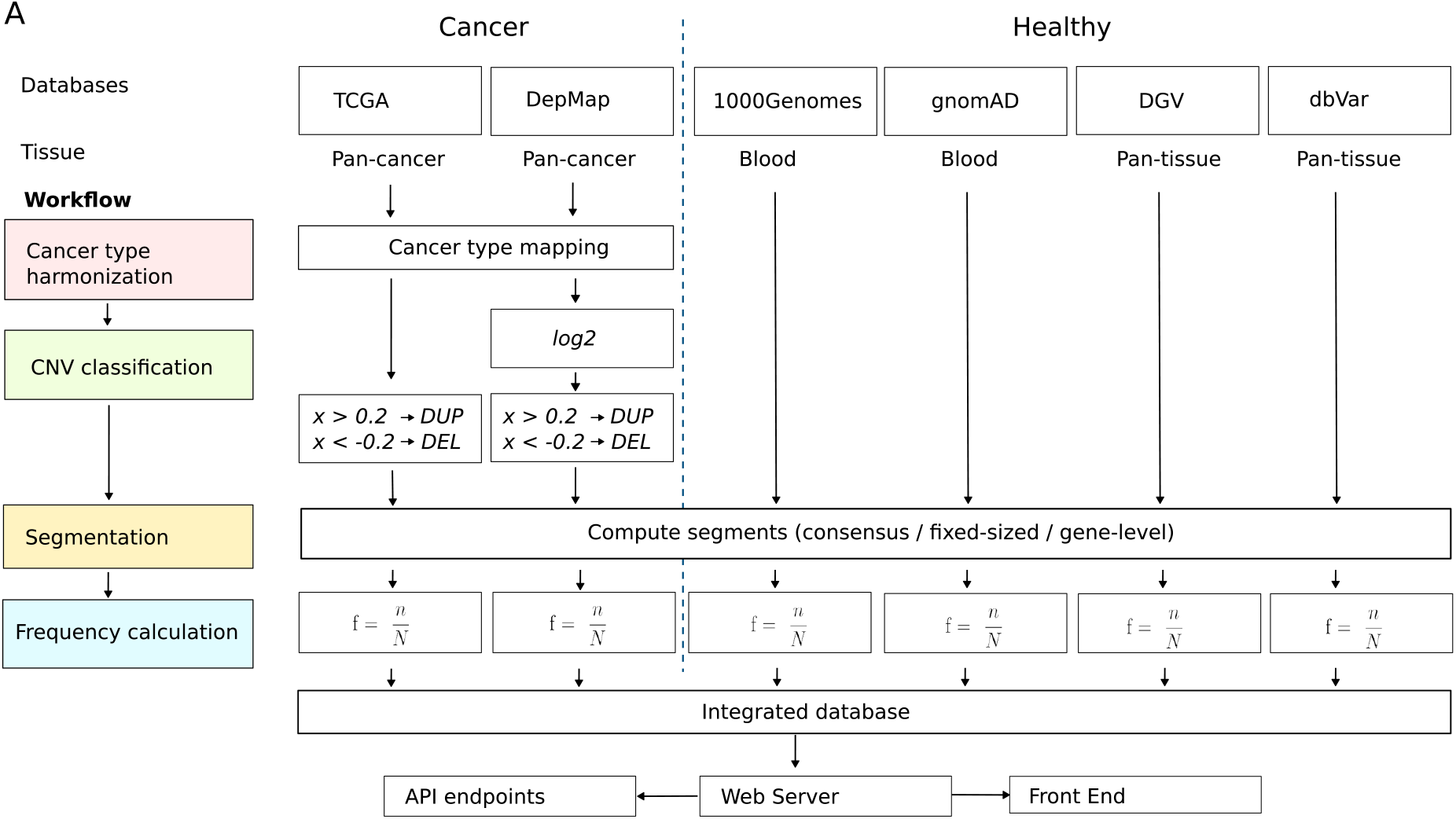
PanCNV-Explorer dataset construction workflow. Segment-level copy number variation (CNV) data were integrated from tumor samples in TCGA and DepMap, alongside germline CNV calls from population databases (1000 Genomes, gnomAD, DGV, and dbVar) to establish baseline genomic variation. After harmonizing cancer type annotations across cohorts, tumor-derived CNVs were classified as amplifications (*log*2 ratio *>* 0.2) ordeletions (*log*2 ratio < −0.2) relative to cohort-specific normal references. To enable multi-scale analysis, three complementary segmentation strategies were applied: 1) consensus segments, 2 fixed-size genomic bins, and 3 gene-level segments. CNV frequencies were then computed per cancer type, CNV type (amplification/deletion), and segmentation strategy.

To enable comprehensive analysis of the CNV data, we created three versions of the database: First, consensus segments were derived by aggregating overlapping CNV calls across all cancer types, followed by hierarchical clustering to define recurrent regions, yielding 200,000 high-confidence segments. Second, fixedsize bins of 10,000 base pairs per segment spanned the entire genome, ensuring uniform resolution for comparative analyses. Third, gene-level segments aligned CNV boundaries to protein-coding gene annotations, facilitating gene-centric interpretation. For each segment, we computed CNV frequencies per mutation type (amplifications/deletions) and per cancer type.

### 3.2 An interactive web resource for pan-cancer CNV analysis

The PanCNV-Explorer web interface enables dynamic exploration of CNV frequencies across genomic regions, cancer types, and CNV classes (somatic/germline). Users can query CNVs by genomic coordinates or by gene symbol, with real-time rendering of amplification/deletion frequencies relative to germline baselines. Users can select one or multiple cancer entities, enabling an interactive comparison across cancer types and normal tissue (Figure 2A). The ability to interactively zoom into the plots, as well as the overlay of known tumor suppressor genes and oncogenes enables a rapid contextualization of CNV events. Additionally, a zoomable genome browser with integrated, dynamically selectable tracks enables a direct comparison of CNV events to transcripts (Figure 2B), regulatory elements, as well as the median gene expression for the selected tissues. To elucidate tissue-specific CNV patterns, UMAP visualization clusters cancer types based on their characteristic CNV frequency profiles (Figure 2C).

**Figure 2.**
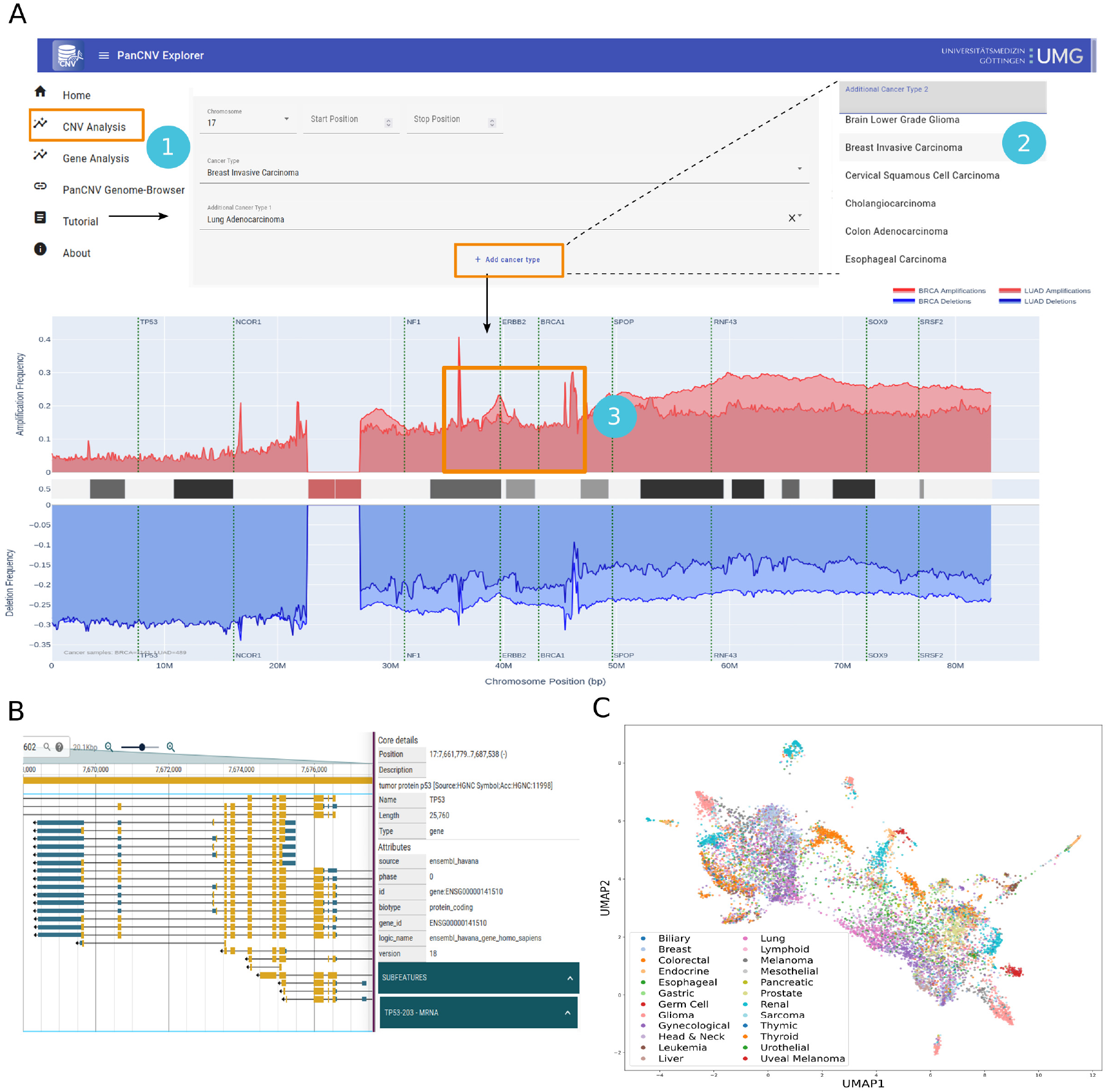
The PanCNV-Explorer web interface enables researchers to investigate cancer-associated copy number variations (CNVs) across tumor types. **(A)** Users can select either 1) a specific genomic region (defined by chromosomal coordinates) or a gene of interest. After choosing a region and 2) one or more cancer types, the tool enables direct comparison of CNV frequency across cancer entities. The CNV frequency visualization plots amplifications (red) and deletions (blue) along the genomic axis. Known tumor suppressor genes (e.g., TP53) and oncogenes (e.g., ERBB2) are marked with dashed lines to highlight their functional roles. By selecting a region within the plot,3) users can interactively zoom in a specific region. **(B)** Aligned genome browser view of the gene transcripts and details, as well as regulatory regions, as **(C)** UMAP visualization of pan-cancer TCGA CNV profiles, revealing strong tissue-of-origin clustering. Distinct clusters for germ cell tumors, renal cell carcinoma, and thyroid cancer highlight cancer-type-specific CNV signatures, consistent with known genomic drivers in these malignancies.

### 3.3 API endpoints and database availability

All analytical modules are accessible via RESTful Application Programming Interfaces (API) end-points, enabling programmatic access to CNV frequency matrices per cancer type and segments, statistical results, and interactive visualizations. The complete database is downloadable in the three formats (consensus, fixed-size bins, gene-level) in CSV format from the PanCNV-Explorer web interface. Additionally, integrated links to the variant interpretation framework Onkopus [41] enable a detailed clinical interpretation of single CNV events.

## 4 Discussion

Copy number variants are pervasive drivers of cancer, yet their systematic analysis in different tissues in cancer types is hindered by fragmented data resources and a lack of integration between cancer-specific alterations and population-level variation. PanCNV-Explorer bridges this gap by presenting a harmonized CNV database consisting of pan-cancer and normal tissue datasets. Coupled with an interactive web interface, it facilitates dynamic exploration of CNV landscapes, from descriptive frequency mapping to mechanistic insights and therapeutic hypothesis generation, while contextualizing cancer-specific alterations against population baselines. Critically, PanCNV-Explorer integrates population germline CNV frequencies as a diagnostic baseline to distinguish cancer-driver events from benign polymorphisms. As breakpoints differ for individual CNVs, a key challenge in CNV data integration is segment harmonization: Overly large segments obscure focal driver events, while excessively granular segments introduce noise from technical artifacts. To resolve this, we developed a consensus segmentation pipeline that merged raw CNV calls, followed by clustering, that provided biologically meaningful segments. This approach effectively captures regions of non-random copy number alteration that likely represent driver events, as it accounts for the natural variation in CNV breakpoints across samples while maintaining sensitivity to both broad chromosomal alterations and focal events. Additionally, we computed pan-cancer CNV frequencies for fixed-sized bins, as well as gene-level frequencies. Unlike GISTIC2.0, which identifies cancer driver peaks in single datasets, PanCNV-Explorer explicitly focuses on pan-cancer analysis, and contrasts somatic frequencies against population baselines to flag events exceeding natural variation (e.g., MYC amplifications). As a limitation, it has to be noted that differences in the CNV data arise due to the different preprocessing of the datasets. On the one hand, the CNVs were called using different software tools, including GATK [12] and CNVnator [42]. All analytical modules—including CNV frequency queries, population-adjusted significance testing, are accessible via RESTful API endpoints, enabling programmatic incorporation into clinical decision support systems or large-scale genomic pipelines.

In summary, PanCNV-Explorer establishes a unified resource for pan-cancer somatic CNVs and population germline variation, with three harmonized data representations and dynamic web/API access. By contextualizing cancer CNVs against population baselines, it transforms observational data into biologically interpretable signals. As cancer genomics transitions toward clinical implementation, resources that rigorously distinguish driver CNVs from noise will be essential for precision oncology. We anticipate PanCNV-Explorer will accelerate biomarker discovery and empower clinicians to interpret CNVs in molecular tumor boards, significantly advancing personalized therapy for cancer patients.

## Data availability

A public instance of PanCNV-Explorer is available at https://mtb.bioinf.med.uni-goettingen.de/pancnv-explorer/. The source code of PanCNV-Explorer, including the database generation, the web server, the web front end and the CNV annotation modules, is available at https://gitlab.gwdg.de/MedBioinf/mtb/cnv-database. The annotation modules for CNVs and structural variants were added to the Onkopus framework and are publicly available through APIs at https://mtb.bioinf.med.uni-goettingen.de/onkopus/api. All data sources of the copy number alteration annotation are publicly available.

## Acknowledgements

The authors thank the members of the molecular tumor board of the University Medical Center Göttingen, especially Alexander König, Li Beißbarth, and Kirsten Reuter-Jessen, for their support and feed-back. The authors thank the International Max Planck Research School for Genome Science (IMPRS-GS).

## Author contributions

JD, NSK and KK conceptualized the methodology and architecture. NSK and ARK constructed the database. NSK, KK and ARK implemented the web server. NSK implemented the annotation modules. NSK and ARK conducted the analysis. NSK, KK and ARK implemented the front end. NSK wrote the manuscript. All authors approved the manuscript.

